# Knowledge.Bio: A Web Application for Exploring, Building and Sharing Webs of Biomedical Relationships Mined from PubMed

**DOI:** 10.1101/055525

**Authors:** Richard Bruskiewich, Kenneth Huellas-Bruskiewicz, Farzin Ahmed, Rajaram Kaliyaperumal, Mark Thompson, Erik Schultes, Kristina M. Hettne, Andrew I. Su, Benjamin M. Good

## Abstract

Knowledge.Bio is a web platform that enhances access and interpretation of knowledge networks extracted from biomedical research literature. The interaction is mediated through a collaborative graphical user interface for building and evaluating maps of concepts and their relationships, alongside associated evidence. In the first release of this platform, conceptual relations are drawn from the Semantic Medline Database and the Implicitome, two compleme
ntary resources derived from text mining of PubMed abstracts.

*Availability*— Knowledge.Bio is hosted at http://knowledge.bio/ and the open source code is available at http://bitbucket.org/sulab/kb1/.

*Contact*—

## I. INTRODUCTION

Knowledge.Bio is an open access web application that provides value-added access to the network of interconnected knowledge latent in the vast scientific literature. Biomedical research scientists, students and the general public can use it to explore relationships between concepts of interest, identifying relevant literature as they develop new hypotheses.

Knowledge.Bio integrates explicit, semantic relations (e.g. stimulates, causes) with statistically predicted, untyped associations between genes and diseases. Currently, the semantic relations are retrieved from the Semantic Medline Database (SemMedDB) [2], a collection of more than 70 million relations mined from PubMed abstracts using the SemRep natural language processing system [3, 4]. The predicted gene-disease associations are incorporated from the “Implicitome”, a resource computed from PubMed abstracts using ‘concept profile’ technology [5].

**Figure 1.**
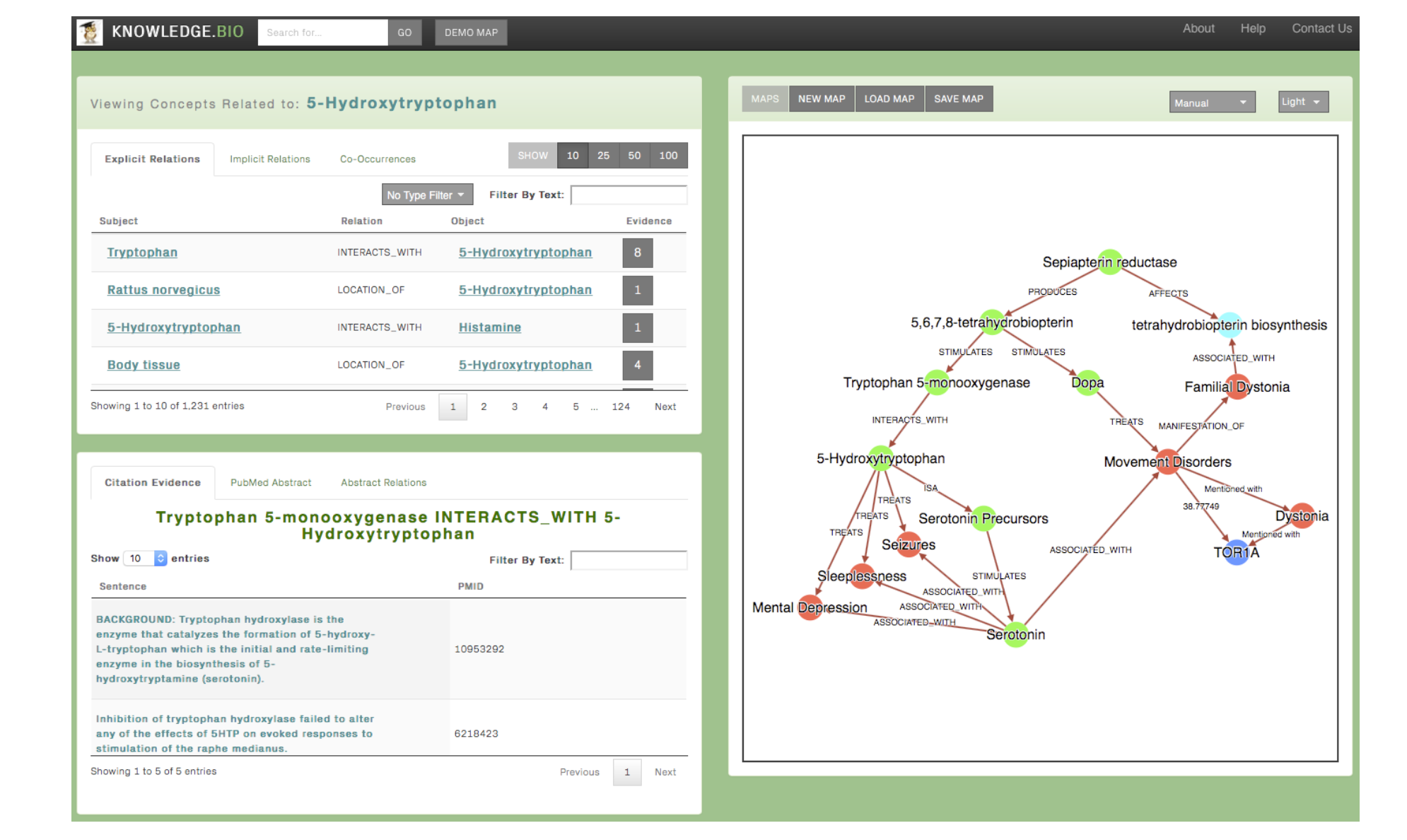
The knowledge. Bio user interface. The upper left panel show relationships linking the 5-Hydroxytryptophan compoundto other concepts such as Histamine. The lower left panel provides evidence for these relations.The map view in the right pane is showing connections between Sepiapterin Reductase (SPR) and 5-Hydroxytryptophan. 5-Hydroxytryptophan was identified as a therapeutic for patients with genetic SPR deficiencies resulting in severe movement disorders [1].discoveries of this nature can be facilitated by the knowledge.bio web application.

**Figure 2.**
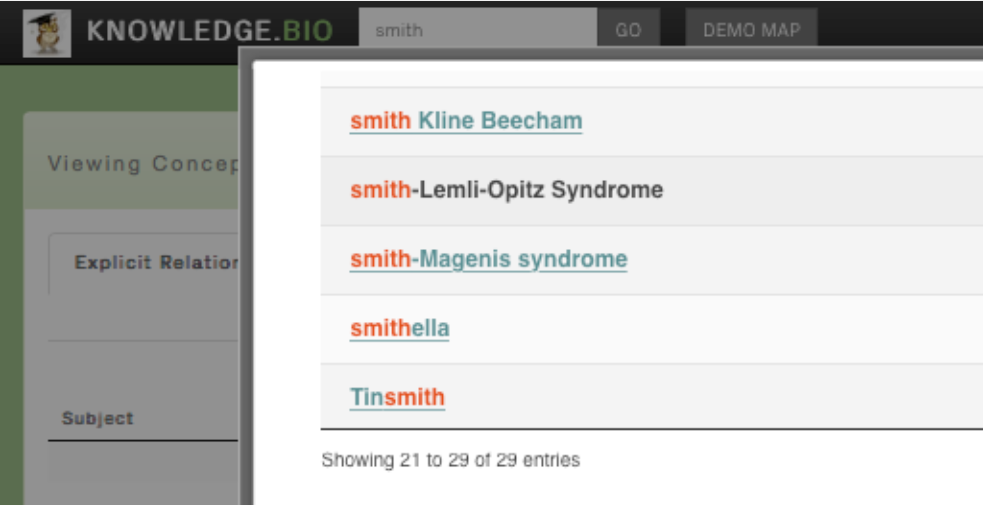
Concept search. Users initiate a session with a text-based search and then select a specific, unique concept to start relation exploration. Here the user has searched for ‘Smith’ and is about to select Smith-Lemli-Opitz Syndrome.

The knowledge in the system is centered on the relationships between uniquely defined concepts, which may either be biomedical concepts from the Unified Medical Language System (UMLS) [6] or genes cataloged within Entrez Gene [7]. The user interacts with the application by searching through this concept space, identifying relations of interest, browsing supporting PubMed references, and iteratively constructing a dynamic network diagram depicting the selected relationships (Figure 1). Below we consider each feature in turn, providing an example of using Knowledge.Bio to identify a new potential gene disease relationship and unearth a hypothetical explanation for the association.

## II. Using Knowledge.Bio

Knowledge.Bio is a “one-stop-shop”, a single page application allowing the user to explore biomedical knowledge.

**Starting a search**. Users initiate exploration by typing-into the search box located to the right hand side of the site logo-some text string expected to match,at least partially,the name of a disease,gene or drug of interest to them. Pressing “Enter”or clicking “Go” results in a popup window highlighting concepts with names matching the search string. Searching for ‘smith’ identifies 29 concepts such as Smith-Lemli-opitz Syndrome that appear in explicit relations (Figure 2).

Selecting one of the enumerated concepts results in retrieval of associated conceptual relationships into tabbed tables on the left hand side of the application: the “Explicit” table presenting relationships retrieved from SemMedDB (Figure 3) and the “Implicit” data table, presenting Implicitome-indexed relationships (Figure 4). When a concept is first selected, a graphical node tagged with the chosen concept name is rendered in the concept map view to the right hand side of the screen.

**Figure 3.**
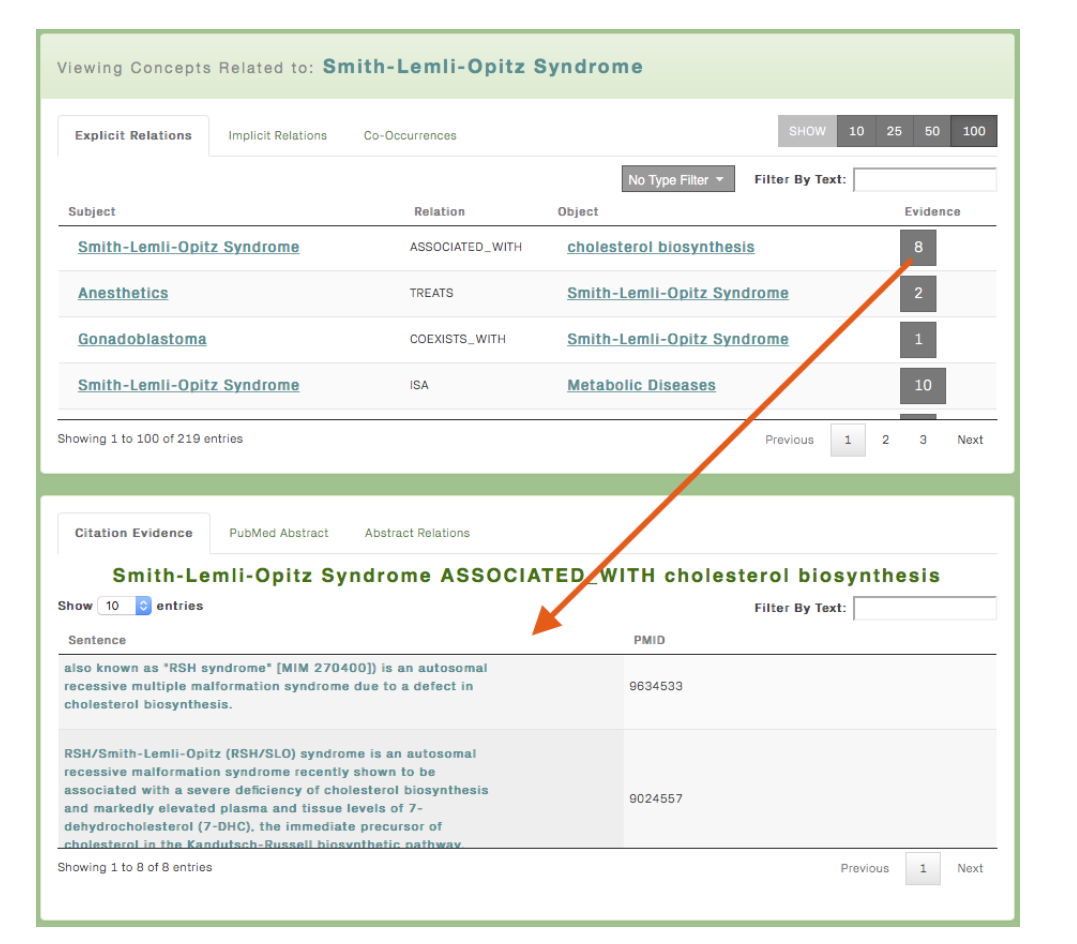
Explicit relations table view. Here, the ‘explicit relations’ tab is selected, displaying 219 relations containing Smith-Lemli-opitz Syndrome as either the subject or or the object. The table may befiltered based on the appearance of specified strings (“filter by text”) or based on the semantic types of the entities in the table (drop down “Type Filter”). The evidence column displays thenumber of PubMed references supporting the predicted relationship. Clicking on the number ofreferences brings up the ‘Citation Evidence’ view displaying the sentences where the relations were found as well as links to the underlying abstract.

**Navigating relation tables**. once a seed concept is selected, the user may continue to iteratively explore the evidence associated with each displayed conceptual relationship. For explicit data relationships, clicking on the “Evidence” button for each row results in the display of matching sentences from PubMed articles in the “Citation Evidence” data table at the lower left hand corner of the screen (Figure 3). Clicking on such citation sentences also displays the associated PubMed abstract. For implicit relationships, clicking on the score field of the relationship displays a “Cooccurrence” list of so-called “B” associated concepts which contribute transitively to the given implicit relationship between theseed “A” concept and matching “C” implicit concept (following Swanson’s ABC model [8] as implemented by the Implicitome ([5]).

**Figure 4.**
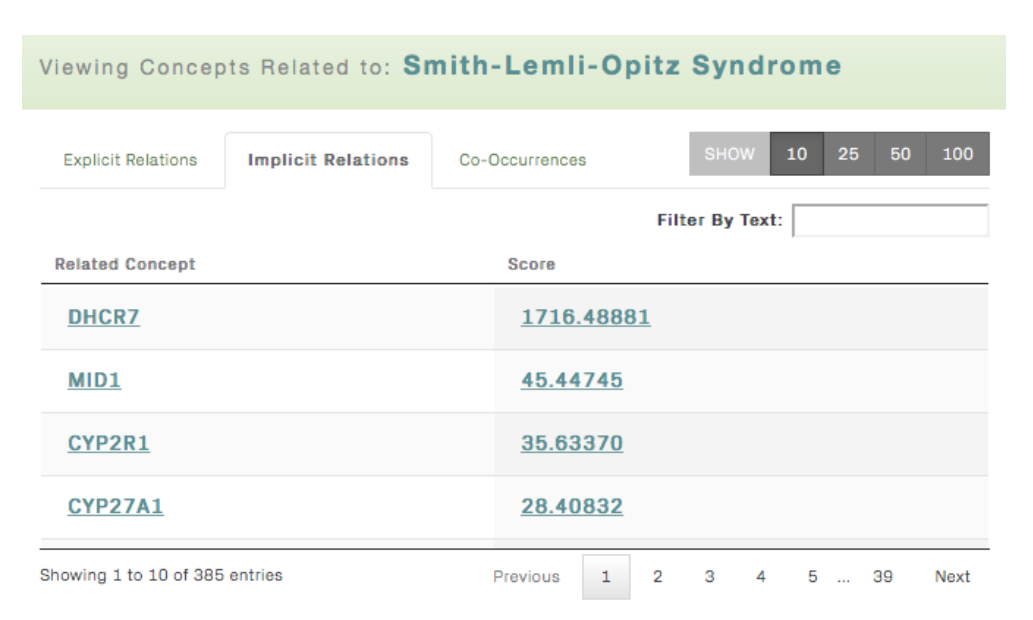
Implicit relations table view. This view shows genes that are predicted to be associated with Smith-Lemli-Opitz Syndrome based on the similarity of their concept profiles.The power of the association is quantified in the ‘Score’column.Clicking on the score will show the key concepts used to connect the gene to the disease in the ‘Co-Occurrences’tab.

In addition, users may click concept links displayed in each of these data tables to bring up a popup window presenting the definition and associated meta-data for the concept. Definitions are drawn either from the UMLS or, in the case of genes, from the mygene.info Web service [9]. The popup window also displays buttons which give the user the option of navigating to that concept’s other relationship data, or to add the concept node and the associated relationship (as an edge) to the user’s concept map. Conceptual relationships may also be added to the concept map by dragging the given row on the left hand data table, over to the right hand concept map graphical view.

**Building concept maps**. As the user navigates the relations provided in the table views, they can manually construct a graphical map of the relations that they deem relevant and trustworthy based on inspection of the provided evidence. This interactive map, rendered in the right hand panel of the interface, provides a way to both visualize and store the network of knowledge identified in their browsing session (Figure 5).Concept labelled graph nodes are color coded by primary concept category:red for disease concepts, blue for gene concepts, green for pharmaceutical concepts including protein products, cyan for pathway concepts, and gray for other concept types such as cell components. Clicking on nodes or edges of the concept map bring up popup windows with node/edge navigation and deletion options.Importantly, the user can reveal the evidence underlyinga relation in the citation view by clicking on the relation edges in the map. As the relations provided by the application are the result of imperfect text-mining, review of the textual evidence underlying each is crucial. For example, some may disagree with the relation in Figure 3 indicating ‘Anesthetics treat Smith-Lemli-Opitz Syndrome’ and thus can choose to remove it from their personal maps. In addition, option widgets located above the concept map view give the user control over “manual” versus various auto-layout map renderings provided by cytoscape.js [10], as well as a “dark versus light” background coloring switch.

**Archiving and sharing concept maps**. The current release of Knowledge.Bio provides buttons to trigger saving and reloading of user-constructed concept maps onto the user's local computer, stored in a Knowledge.Bio-specific formatted text file. The text file contains tab delimited representations of each relation in the map followed by a JSON object that stores the layout. A collection of sample Knowledge.Bio maps that can be loaded into the application can be found at https://figshare.com/articles/Demonstration_knowledge_bio_graphs/3394114.

**Generating a hypothesis**. The map displayed in Figure 5 encapsulates one example discovery process that could lead to a testable hypothesis (also described in [5]). Beginning with a search for Smith-Lemli-Opitz Syndrome,the user first selects the implicit relations tab and identifies CYP2R1 as potentially relevant gene (Figure 5A). Next,the user identifies DHCR7 as a known causative gene based on multiple explicit statements in the literature (Figure 5B).Finally,the user identifies explicit statements indicating that both DHCR7 and CYP2R1 affect Vitamin D metabolism (Figure 5C,D), suggesting this as a potentially common mechanism linking CYP2R1 to Smith-Lemli-Opitz Syndrome.

## III. Discussion

Since Swanson’s introduction of the paradigm of ‘literature based discovery’ (LBD) [8], a variety of LBD tools have been introduced. Many of the these are reviewed in [11, 12] though most of the approaches mentioned are either not represented in user friendly applications or are no longer publicly available. A few examples of tools that are online at the time of this writing include Semantic MEDLINE [13], EpiphaNet [14], and Arrowsmith [15]. Semantic MEDLINE and EpiphaNet both expose semantic relationship data from SemmedDB, rendering graphical networks in responseto user queries. EpiphaNet extends the relations in Semantic MEDLINE with relation rankings and predicted associations based on distributional semantics, a methodology that, as in the Implicitome, quantifies the likelihood that two concepts are related based on the similarity of their vectors of co-occurring concepts. EpiphaNet’s current online form (http://epiphanet.uth.tmc.edu), varies from that described in the original manuscript in that focuses exclusively on identifying links between drugs and side effects whereas Knowledge.Bio and Semantic MEDLINE span interconnections between more than 50 different semantic types. Semantic MEDLINE focuses on establishing a process through which the user can rapidly iterate with the interface as they explore the new relationships that it brings to their attention - a process that they refer to as “cooperative reciprocity”. Arrowsmith lacks a network-based visualization, it focuses only on identifying the most relevant intervening ‘B’terms linking two input “A” and“C” terms.

**Figure 5.**
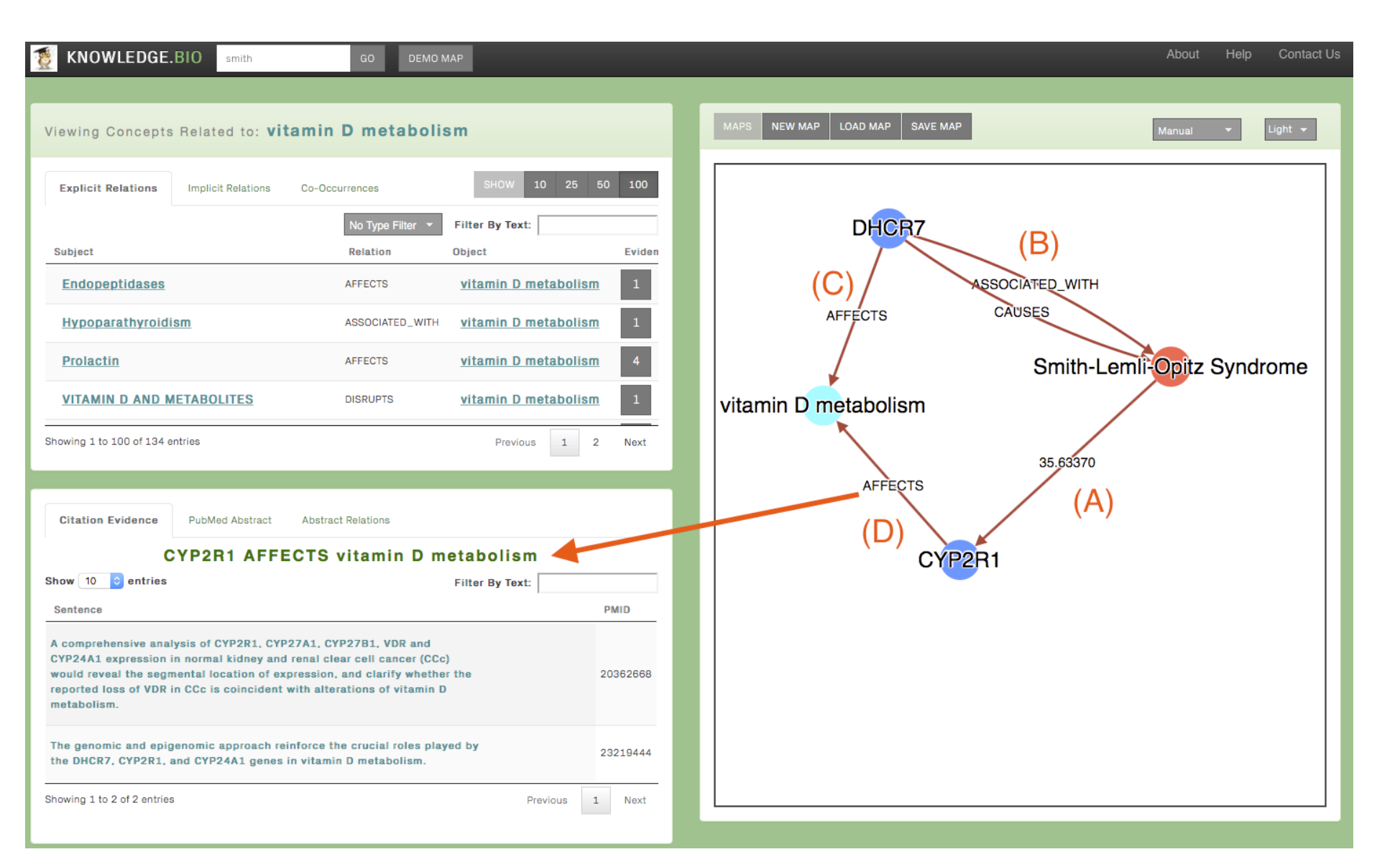
Knowledge.bio map. The map shows an implicit connection between Smith-Lemli-Opitz Syndrome and the gene CYP2R1 (A). Further, it shows that CYP2R1 shares a relationship to Vitamin D metabolism with DHCR7 (C,D), a gene known to be genetically associated with Smith-Lemli-Opitz Syndrome (B). Selecting the edges in the map brings up the evidence underlying the relation (as indicated by the red arrow).

Knowledge.Bio is similar in process to Semantic MEDLINE in that it relies on the user to assess the relevancy of the SemRep-mined relations it exposes to them. Key differences are: (1) Knowledge.Bio integrates the implicit gene-disease relations calculated by the Implicitome,(2) the Knowledge.Bio graph is entirely user-created and can be saved and shared where the Semantic MEDLINE graph is an automatically generated view, (3) Semantic MEDLINE requires a UMLS account while no login is needed for Knowledge.Bio, (4) Semantic MEDLINE sessions are oriented around free-text searches into PubMed with semantic predictions drawn from the corresponding abstracts while Knowledge.Bio search is oriented around individual unique concepts.

In future versions of Knowledge.Bio, we hope to build on some of the algorithmic advances of EpiphaNet and related work to provide the user with optional enhanced guidance in their explorations. In particular, we hope to extend the current single-concept search with smart pathfinding options to support “closed discovery” (e.g. given A and D, identify meaningful A->B…C->D paths that relate them. Apart from these pathfinding services, the application will be extended to incorporate additional 3rd-party data sources such as Gene Ontology annotations that will substantially extend the interconnectedness of its underlying knowledge graph. Finally, and perhaps most importantly, the new version will provide facilities for storing, sharing, discussing and online archiving of user-contributed and user-verified conceptual relationships. These social, collaborative features are intended to stimulate the evolution of the Knowledge.Bio knowledge base both at the level of individual research projects and at the more global level, weeding out incorrect or uninteresting assertions and expanding the system based on the user community's collective knowledge.

## IV. Conclusion

Knowledge.bio provides a novel Web interface for user discovery and interpretation of research literature,indexed in personal concept maps.The application is responsive, stable, requires no authentication,and provides empowers its users with otherwise hard-to-access text-mined information in a simple, user-friendly manner.Further, the application is completely opensource and could easily be extended or adapted by motivated developers.

**Implementation**. Knowledge.Bio is available online at the web site http://knowledge.bio, with open source code available at http://bitbucket.org/sulab/kb1. It consists of a Python Django web application framework enhanced with custom JavaScript code leveraging various general purpose web client libraries,in particular, Jade templates (http://jade-lang.com/), Bootstrap.js (http://getbootstrap.com) and, DataTable.js(http://www.datatables.net).The right hand sideconcept map graph view is rendered using cytoscape.js (Franz *et al.* 2016).The back end is a MySQL database schema derived from published SemmedDB, UMLS and Implicitome data models.

## Acknowledgements

We would like to thank P.A.C. 't Hoen and Marco Roos from the Leiden University Medical Center for testing the software.

## Funding

This work was supported by the US National Institute of Health (grants GM089820 and U54GM114833 to AIS) and by the Scripps Translational Science Institute with an NIHNCATS Clinical and Translational Science Award (CTSA; 5 UL1 TR001114). Funding from the European Commission (FP-7 project RD-Connect, grant agreement No. 305444), and ODEX4all (NWO 650.002.002) are gratefully acknowledged.

## Conflict of Interest

Kristina M. Hettne has performed paid consultancy since November 1, 2015, for Euretos b.v, a startup founded in 2012 that develops knowledge management and discovery services for the life sciences, with the Euretos Knowledge Platform as a marketed product. The paid consultancy did not specifically fund this study. Richard M. Bruskiewich is CEO of STAR Informatics/Delphinai Corporation, the Canadian firm which provides software engineering services for this project under subcontract to Scripps using funds from the Scripps-hosted research grants.

## References

1. Bainbridge MN, Wiszniewski W, Murdock DR, FriedmanJ, Gonzaga-Jauregui C, Newsham I, Reid JG, Fink JK, Morgan MB, Gingras MC et al: Whole-genome sequencing for optimized patient management. Sci Transl Med 2011, 3(87):87re83.

2. Kilicoglu H, Shin D, Fiszman M, Rosemblat G,Rindflesch TC: SemMedDB: a PubMed-scale repository of biomedical semantic predications. Bioinformatics 2012, 28(23):3158–3160.

3. Rindflesch TC, Fiszman M: The interaction of domain knowledge and linguistic structure in natural language processing: interpreting hypernymic propositions in biomedical text. J Biomed Inform2003, 36(6):462–477.

4. Ahlers CB, Fiszman M, Demner-Fushman D, Lang FM,Rindflesch TC: Extracting semantic predications from Medline citations for pharmacogenomics. Pac Symp Biocomput 2007:209-–220.

5. Hettne KM, Thompson M, Van HaagenHH, van derHorst E, Kaliyaperumal R, MinaE, TatumZ, Laros JF, van Mulligen EM, Schuemie M et al: The Implicitome: A Resource for Rationalizing Gene-Disease Associations. PLoS One 2016,11(2):e0149621.

6. Bodenreider O: The Unified Medical Language System (UMLS): integrating biomedical terminology. Nucleic acids research 2004, 32:267-270

7. Maglott D, Ostell J, Pruitt KD, Tatusova T: Entrez Gene: gene-centered information at NCBI. Nucleic acids research 2011, 39:D52–57.

8. Swanson DR: Fish oil, Raynaud's syndrome, and undiscovered public knowledge. Perspectives in biology and medicine 1986, 30:7–18

9. Wu C, Macleod I, Su AI: BioGPS and MyGene.info: organizing online, gene-centric information. Nucleic Acids Res 2013, 41 (Database issue):D561–565.

10. FranzM, Lopes CT, HuckG, Dong Y, Sumer O, Bader GD: Cytoscape.js: a graph theory library for visualisation and analysis. Bioinformatics 2016, 32 (2):309–311.

11. Jensen LJ, Saric J, Bork P: Literature mining for the biologist: from information retrieval to biological discovery. Nat Rev Genet 2006, 7(2):119– 129.

12. Weeber M, Kors JA, Mons B: Online tools to support literature-based discovery in the life sciences. Brief Bioinform 2005, 6 (3):277–286.

13. Cairelli MJ, Miller CM, Fiszman M, Workman TE, Rindflesch TC: Semantic MEDLINE for discovery browsing: using semantic predications and the literature-based discovery paradigm to elucidate a mechanism for the obesity paradox. AMIA Annu Symp Proc 2013, 2013:164–173.

14. Cohen T, Whitfield GK, Schvaneveldt RW, Mukund K, Rindflesch T: EpiphaNet: An Interactive Tool to Support Biomedical Discoveries. J Biomed Discov Collab 2010, 5:21–49.

15. Swanson D: An interactive system for finding complementary literatures: a stimulus to scientific discovery. Artificial Intelligence 1997, 91:183–203.

